# Only ten percent of the global terrestrial protected area network is connected via intact land

**DOI:** 10.1101/2020.01.28.920488

**Authors:** Michelle Ward, Santiago Saura, Brooke Williams, Juan Pablo Ramírez-Delgado, Nur Arafeh-Dalmau, James R. Allan, Oscar Venter, Grégoire Dubois, James E.M. Watson

**Affiliations:** School of Earth and Environmental Sciences, University of Queensland, St Lucia 4072, Australia; Centre for Biodiversity and Conservation Science, School of Biological Sciences, The University of Queensland, Brisbane 4072, Queensland, Australia; European Commission, Joint Research Centre (JRC), Via E. Fermi 2749, I-21027 Ispra, VA, Italy; ETSI Montes, Forestal y del Medio Natural, Universidad Politécnica de Madrid, Ciudad Universitaria s/n, 28040, Madrid, Spain; Natural Resource and Environmental Studies Institute, University of Northern British Columbia, Prince George, Canada; Institute for Biodiversity and Ecosystem Dynamics (IBED), University of Amsterdam, P.O. Box 94240, 1090 GE, Amsterdam, The Netherlands; Wildlife Conservation Society, Global Conservation Program, Bronx, NY 20460, USA

## Abstract

Land free of direct anthropogenic disturbance is considered essential for achieving biodiversity conservation outcomes but is rapidly eroding. In response, many nations are increasing their protected area estates but little consideration is given to the context of the surrounding landscape. This is despite the fact that connectivity between protected areas is critical in a changing climate and mandated in international protected area targets. By utilizing the latest human pressure assessment, which shows that ∼40% of the terrestrial planet is still intact, and a connectivity method, we found just 9.7% of Earth’s terrestrial protected network can be considered ‘connected’. On average, 11% of each nation’s protected area estate is connected via intact land. As the global community commits to bolder action on abating biodiversity loss, only an increased focus on landscape-scale habitat retention and restoration efforts will ensure those critical areas safeguarded for conservation outcomes will remain (or become) connected.

**One Sentence Summary:** Only 9.7% of the global protected area network can be considered ‘connected’ by intact land.

## Main Text

Protected areas (PAs) are a core tool in abating the biodiversity crisis (*1*), and their importance is reflected in the 2020 Strategic Plan for Biodiversity (*2*). This international agreement calls for the expansion of the global PA network to cover 17% of terrestrial areas and 10% of marine areas by 2020. Crucially, the Strategic Plan stipulates that PA networks must be well-connected, effectively and equitably managed, and also cover ecologically representative areas of particular importance for biodiversity (*2*). Due to their extraordinary importance for biodiversity outcomes, PAs have received substantial research attention in global conservation policy discussions, with recent assessments focusing on how well they are representing species (*3*), their overall management effectiveness (*4*), and how well are they abating key threatening processes (*5*). Yet, to date, reporting has been almost completely blind to how well connected the expanding global PA estate is, with the only substantive research considering connectivity solely through protected land (*6*), disregarding the condition of the wider landscape context.

High levels of landscape connectivity is seen as critical for species adaptation under anthropogenic climate change as it allows individuals and populations to track their preferred climatic conditions (*7*). Under projected climate scenarios, it is predicted that many species will need to move further and more rapidly in the 21^st^ century and connected landscapes that facilitate this movement is one of the best conservation responses (*8, 9*). Connected landscapes allow fundamental ecological mechanisms to operate unimpeded, such as meta-population retention (*10*) and successful dispersal and migration (*11*). Landscape connectivity allows for increased ecosystem function and resilience (*12*) by ensuring nutrient cycling can continue unabated, as well as other important abiotic conditions such as radiation, wind, light regimes, humidity, and key hydrological regimes (*13, 14*). It is well known that land use activities such as farming, urbanization, mining, and unsustainable forestry disrupt the connectivity of landscapes to various degrees (*11*). The retention, and where necessary, restoration of connectivity across a landscape matrix between protected areas (PAs) is therefore vital for achieving biodiversity goals clearly outlined in the Strategic Plan for Biodiversity (*2*).

Here, we analyze the connectivity of the global terrestrial PA system using measures of contiguity of intact land (areas largely devoid of high anthropogenic pressures that significantly alter natural habitat) as a favorable movement pathway between PAs. We determined the connectivity of the global network of PAs by quantifying intact continuous pathways between PAs. To map intactness, we used the human footprint dataset (HFP), which, at a 1km^2^ resolution, is the most comprehensive, fine-scale human pressure map available as it takes into account agricultural lands, roads, railways, human population density, built environments, night-time lights, and navigable waterways, all of which are driving the species extinction crisis (*15*). We used an updated version of the HFP updated to 2013, which supersedes the published version dated for 2009 (*16*). Following other studies (*5, 17*), we defined intact land as that with a human footprint value <4 out of 50, as this threshold is where anthropogenic activities have significantly changed the state of land from largely natural in extent to highly modified (*18*). Moreover, this human pressure threshold is associated with the sharpest declines in mammal movement (*11*) and is one of the strongest predictors of mammal species extinction risk (*19*). For these reasons, we argue that areas with human pressure above this threshold are unlikely to hold sufficient connectivity value for many key elements of biodiversity. However, recognizing that there is no one true level of pressure that prohibits connectivity for all biodiversity (as ecological responses to human pressure are idiosyncratic), we conducted several sensitivity analyses around this human footprint threshold and found our results were robust to lower and higher human footprint values (table S1 and table S2).

Globally, while 41.6% of terrestrial land is intact, only 9.7% of the area under protection can be considered connected through intact landscapes (Fig. 1 and fig. S1). This means that very few PAs have a fully continuous pathway through intact lands, connecting their demarcated edges. At a continental scale, PAs in Oceania are the most connected across all continents (16.7%), followed by the Americas (14.8%). In contrast, Asia (3.3%), Africa (0.4%), and Europe (0.3%) have extremely low levels of PA connectivity provided by intact lands.

**Fig. 1.**
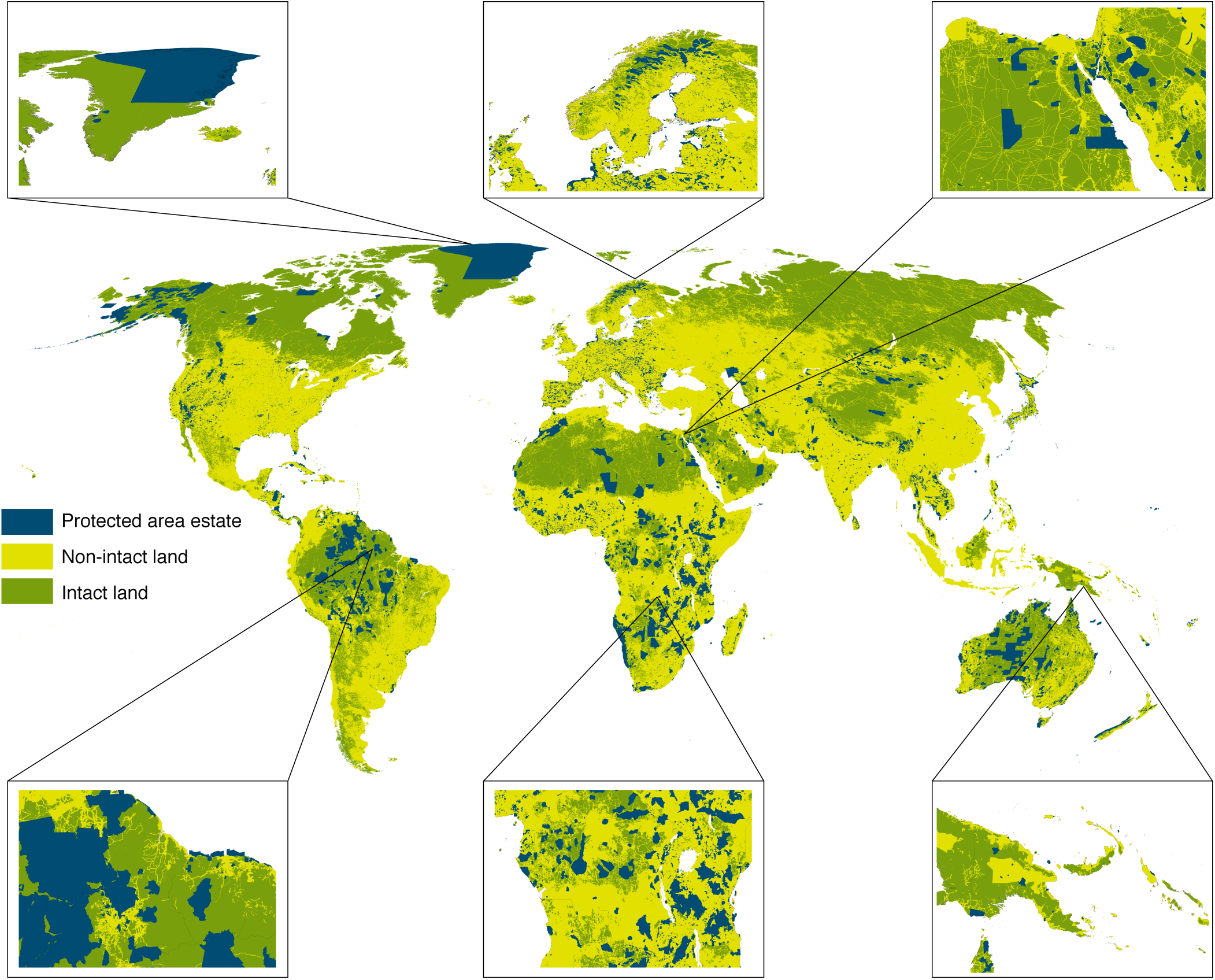
Human pressure compromises protected area connectivity. The spatial distribution of the protected area estate (blue) littered throughout non-intact (yellow) and intact land (green). We provide six fine-scale examples, starting top left and moving clockwise, Greenland, Finland, Egypt, Papua New Guinea, Democratic republic of Congo, and Guyana.

At a national scale, the percentage of connected PAs varies enormously (Fig. 2A and fig. S2 to fig. S5). The majority of nations maintain the lowest level of connectivity possible (median connectivity = 0%). These countries are found not just concentrated in Europe (where one would expect minimal presence of low human pressure matrix between PAs) but also across Asia and Africa (table S3) where landscapes are rapidly changing through large-scale infrastructure projects such as roads and agriculture (*20, 21*). This has significant ramifications for international conservation agendas, as many Asian and African countries are megadiverse when it comes to biodiversity (*22, 23*). For example, Vietnam – one of Earth’s most biologically diverse countries (*24*), has ∼8% protection and no connected PAs based on our analysis, (Fig. 2B) yet PA connectivity is likely vital for the persistence of critically endangered species such as saola (*Pseudoryx nghetinhensis*) and Indochinese tiger (*Panthera tigris corbetti*) (*25, 26*). Similarly, Madagascar is home to some of the most genetically-diverse species on Earth, including the black and white ruffed lemur *(Varecia variegata)*, aye-aye *(Daubentonia madagascariensis)*, and fossa *(Crytoprocta forex)*. These genetically-unique species, being predominately arboreal, require contiguous intact landscapes to fulfill their important ecological roles (*27*), yet we found Madagascar only has 4.2% intact land remaining and no fully connected PAs.

**Fig. 2.**
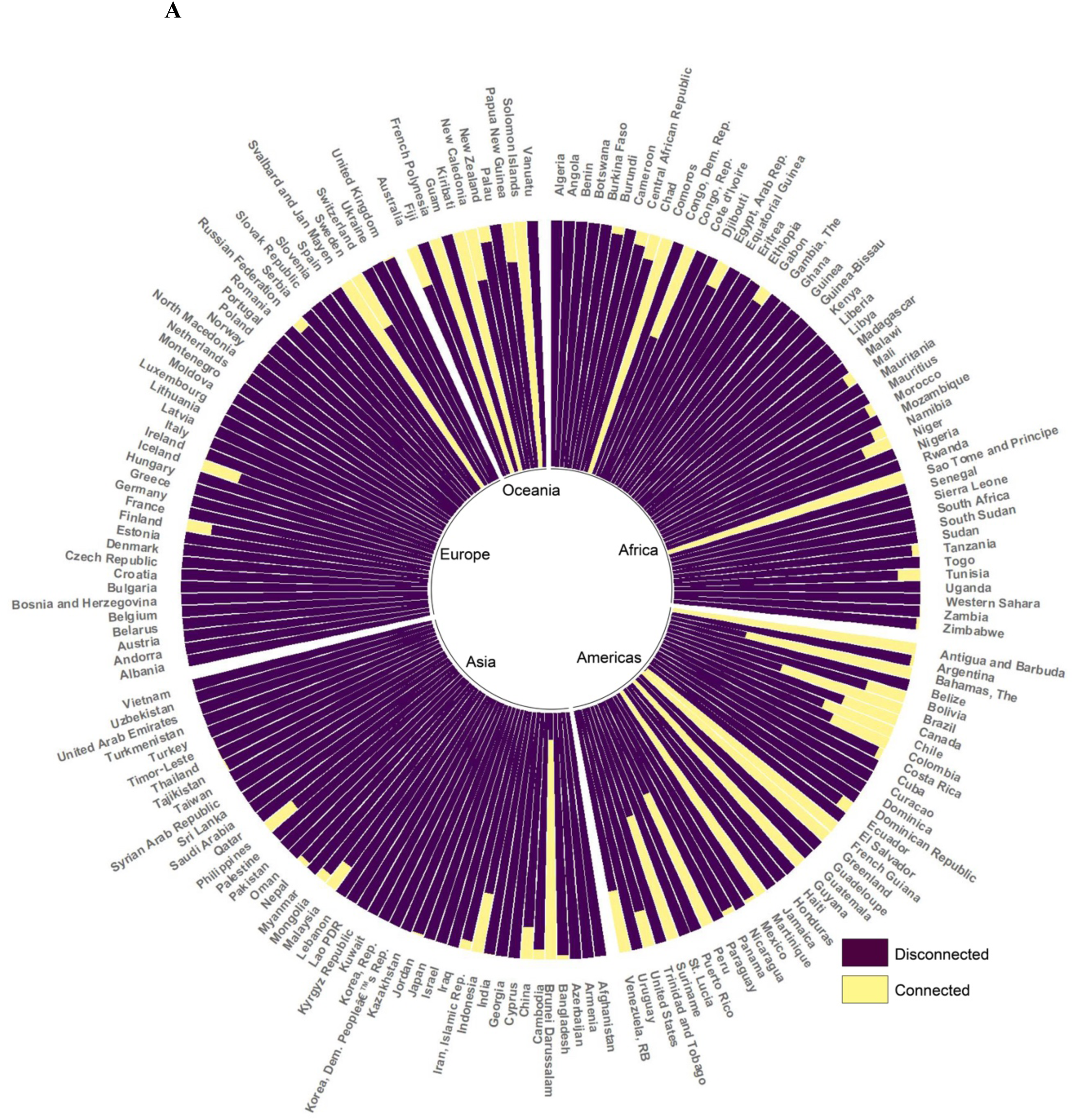
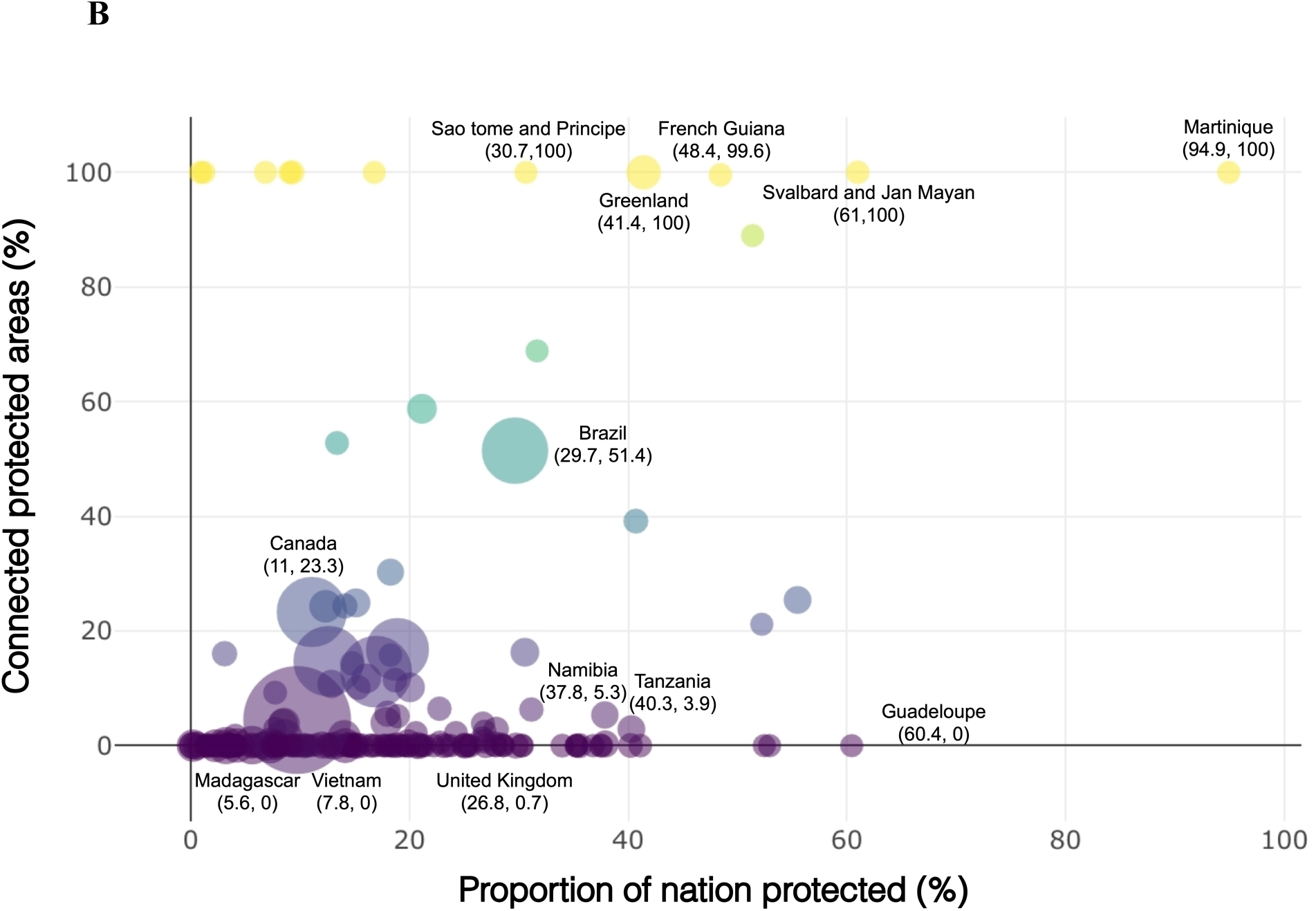
Few nations maintain a connected protected network through intact land. **(A)** Proportion of connected and disconnected protected areas for each nation. Countries are grouped per continent. **(B)** The relationship between the proportion of land under protection (x-axis) and the proportion of connected protected areas (y-axis) per country (less connected countries are purple while more connected countries are yellow). The size of the bubble indicates the size of the country.

Only nine (4.6%) countries have >17% of their land protected (which many nations define as their contributions to the CDB PA target) (*28*) and maintain >50% connectivity across their PA network (Fig. 3). These countries with high proportions of land under protection have statistically significant correlations with higher proportions of connected PAs (ρ=0.28, n=183, p<0.05). We found no significant relationship between countries with high proportions of the protected land connected and the number of PAs (ρ=0.00, n=183, p>0.05) or the size of the country (ρ=0.04, n=183, p>0.05). This indicates that connectivity may not be considered when nations are adding to their PA estates. For the more intact countries, it may be because they declare fewer but larger PAs, while more human-modified countries (such as those in Europe) declare many small PAs that are usually surrounded by a non-intact matrix.

**Fig. 3.**
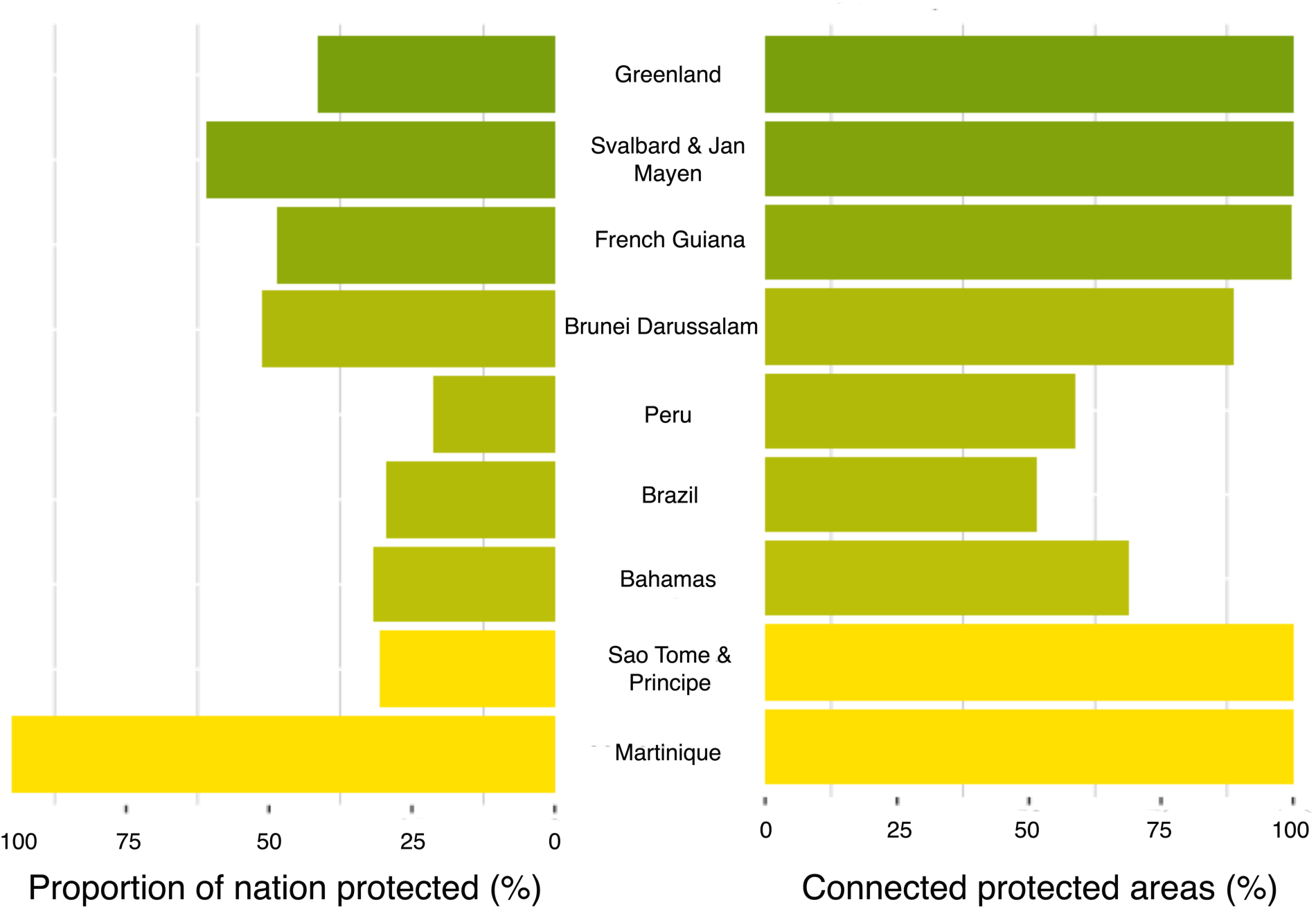
Examples of countries with high proportions of land protected and connected protected areas. Only nine countries protect >17% of land (left) and maintain >50% connected protected areas (right). Countries are ordered based on the highest proportion of remaining intact land (less intact countries are yellow, while more intact countries are green).

Eighteen countries have >50% of their land that can be considered intact, yet have very low connectivity between their PAs (Fig. 4 and fig. S6 to fig. S7). For example, our analysis revealed that Egypt has 77.1% intact land, but only 10.7% of its PA network is connected. Even though there are large PAs within this nation, including El-Gelf El-Keber (48,523km^2^) and Elba (36,600km^2^), major roads, agriculture, and urban sprawl are increasingly fragmenting the landscapes (*29*).

**Fig 4.**
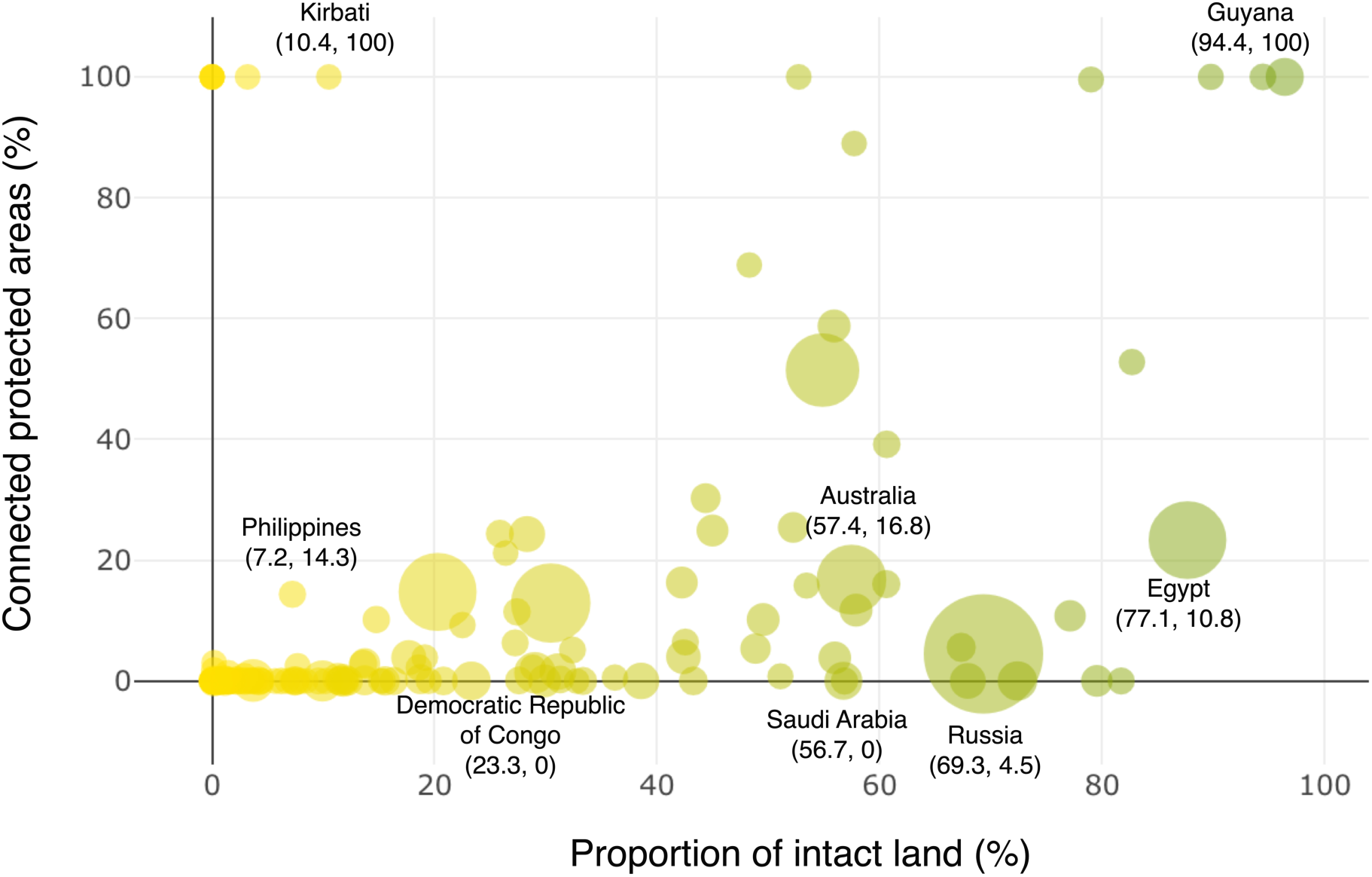
The relationship between the proportion of land that can be considered intact per country and the proportion of connected protected areas. Scatterplot showing the relationship between proportion of intact land (less intact countries are yellow, while more intact countries are green) and proportion of connected protected areas per country. The size of the bubble indicates the size of the country.

The vast majority (76.4%) of countries have very few intact landscapes remaining and low proportions of connected PAs. These include surprising examples such as the Democratic Republic of Congo (DRC), which we found to have 23% intact land and no connected PAs. Much of the PA network within the DRC is scattered across the country, disconnected by agricultural lands, urban areas, and mines. In addition to the 149 countries that have low proportions of both intact landscapes and few connected PAs, there are also 13 countries that do not have any remaining intact landscapes and therefore no connectivity between PAs. Statistically, countries with low proportions of intact land also have significantly lower connected PAs (Spearman’s p=0.40, n=183, p<0.05; Eigenvalue 1.37, Eigenvalue 0.62).

Our results show that while many countries are meeting the areal component of global PA targets, much of this PA estate is not connected through non-modified habitat pathways. This suggests that the overall strategic goal of overcoming biodiversity loss will be likely compromised without increased focused on wider land-use efforts to retain and restore habitats beyond PA boundaries. Any plan for maintaining and restoring ecosystem connectivity between protected areas must include a clear focus on retention strategies for the remaining ecosystems that are currently not degraded because these places are, by definition, areas of high connectivity and known to hold exceptional value (*11, 30*). It is also well accepted that avoiding degradation of habitat (and hence loss of connectivity) is a far better strategy than attempting restoration after it is lost. Restoration is more costly, riskier, and unlikely to lead to a full recovery of connectivity values (*31*). For Earth’s remaining intact, connected ecosystems to be retained, they must be formally recognized, prioritized in spatial plans, then placed under effective management so they are protected from human impacts. Opportunities to recognize and conserve intact lands that help retain connectivity are expanding beyond areas in strict protected areas. For example, a definition for ‘other effective area-based conservation measures’ (OECMs) was agreed by nations in November 2018, with the objective to “…achieve positive and sustained long-term outcomes for the in-situ conservation of biodiversity, with associated ecosystem functions and services and where applicable, cultural, spiritual, socio–economic, and other locally relevant values” (*32*). The global extent of OECMs is expected to increase rapidly over the coming years (*33*). As such, their use, if planned and implemented well, could play an enormous role in keeping connectivity between PAs.

But our findings show that even if humanity was successful in halting the degradation of all remaining intact ecosystems, there would be many PAs isolated given their surrounding matrix has already been highly altered. As such, there is also the need for a broad, transboundary restoration agenda to rapidly increase connectivity between those lands set aside for conservation. There is clearly appetite for such bold restoration action, with the UN recently declaring the ‘Decade on Ecosystem Restoration’ (*34*), by which 350 million hectares of degraded land will be restored between now and 2030. We argue that these types of restoration goals should be framed within a broader connectivity agenda and specifically planned to maximize the quality of the landscape matrix between those PAs essential to biodiversity outcomes. As restoration of less degraded natural ecosystems presents some of the most cost-effective restoration opportunities (*35*) and lowest potential for conflicts with other priorities (such as agriculture), it is essential to incorporate the cost of such restoration to other societal goals when planning for restoration action.

Our results highlight the need for a far more comprehensive reporting framework on area-based conservation that captures not just the extent and overall effectiveness of local implementation of conservation activities but also how connected these PA networks are within the wider landscape. As the relationship between the placement of area-based conservation activities, how they are managed, and the wider landscape context is nuanced, simple metrics on connectivity must be integrated with other assessments of PA effectiveness. For example, while Venezuela is currently achieving 55.5% protection, we found very little connectivity between PAs. This may not be a poor result, as the PA estate within Venezuela seem to be extremely large, well-managed, and representative (*36*). Other nations may have low proportional connectivity because their PAs have been established in human-dominated landscapes, which are often where much conservation action must occur. This type of reactive conservation strategy is clearly necessary for countries that have degraded, but ecologically important, landscapes (*37*). In contrast, some nations that have good connectivity scores could be hiding inherent biases in PA placement. For example, Australia is achieving ∼17% connectivity via our assessment and has ∼19% of its land under protection, which could be considered a relatively good outcome. Yet upon closer inspection, most new PAs have been placed in the desert ecosystems, which do not adequately represent all taxonomic groups and are already over-represented in the nation’s PA estate (*38*). In addition, most Australian PAs are also not effectively managed, with 1,390 threatened species continuing to lose critical threatened species habitat inside and outside PAs (*39, 40*). The numbers around PA connectivity alone can hide important issues but there are now methodologies that capture quality, extent, and connectedness of landscapes (*17*) and it should be possible for nations to transform these into a framework that holistically assesses the overall effectiveness of area-based conservation action.

We note that in some cases, a low proportional connectivity score between PAs may be a reasonable outcome. This is likely the case in some parts of Europe, where small-scale efforts to create corridors between PAs may not be captured at the 1km^2^ resolution of our analysis. In addition, areas of high pressure in the matrix between PAs may be acceptable for some species that have co-existed with human-modified landscapes for centuries. But as most species, especially those that are endangered, cannot persist in human-dominated landscapes (*11, 19*), our results provide a sobering assessment.

Well managed PAs and retaining and restoring intact landscapes that surround them are critical to abating the biodiversity crisis. Right now, the majority of PAs are isolated by a matrix of rapidly eroding intact habitat and are unlikely to be effective, especially in a time of anthropogenic climate change. Our results show that urgent change in how nations protect and restore landscape-scale habitat is crucial as the international community gears up to embrace a new global biodiversity framework post-2020.

## Materials and Methods

We determined the connectivity of the global network of protected areas by quantifying intact continuous pathways (areas largely devoid of high anthropogenic pressures) between protected areas.

### Protected areas

Data on protected area location and boundary were obtained from the May 2019 World Database of Protected Areas (WDPA) (*41*). We only considered protected areas that had a land area of at least 10 km^2^. As China removed most of its protected areas from the public May 2018 WDPA version, we used the April 2018 WDPA for China only, which contained the full set of Chinese protected areas at the time. It is important to note that our statistics may differ from those reported by countries due methodologies and datasets differences used to measure terrestrial area of a country or territory.

### Measure of human pressure

We used the latest global terrestrial human footprint maps – a cumulative index of eight variables measuring human pressure on the global environment – to calculate the average human pressure between protected areas. The Human Footprint 2013 map uses the following variables: 1) the extent of built human environments, 2) population density, 3) electric infrastructure, 4) crop lands, 5) pasture lands, 6) roads, 7) railways, and 8) navigable waterways. Each human pressure is scaled from 0-10, then weighted within that range according to estimates of their relative levels of human pressure following Sanderson et al., 2012. The resulting standardized pressures were then summed together to create the standardized Human Footprint maps for all non-Antarctic land areas (*16*).

Within the main manuscript, we defined intact land as any 1km^2^ pixel with a human footprint value not higher than 4, not including the value 4. Within this threshold, all areas with a human footprint score higher than 4 are defined as non-intact. While previous analyses showed that a >4 score is a key threshold above which species extinction risk greatly increases (*19*), we recognized that there is no one true threshold which impacts all species equally. Some species may require no human pressure to successfully disperse, while others might successfully navigate through more intensively modified landscapes. Therefore, we conducted out analyses for two additional human footprint thresholds. The first used a human footprint score <1 and the second incorporated all areas with a human footprint <10.

### Probability of Connectivity

The Probability of Connectivity network-based metric underlies the analysis performed (*6*), with adaptations to account for connectivity provided by intact lands. Probability of Connectivity is given by the following formula:

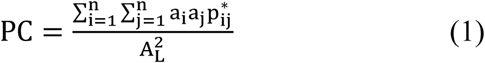

where n is the total number of protected areas (PAs) in the study area (i.e. continent or country), a_i_ is the total area of PA i, p*_ij_ is the maximum product probability between protected areas i and j, and A_L_ is the total area of the study area. The maximum product probability (p*_ij_) considers both direct connections (movement from i to j without using any other intermediate PA in the network) and indirect connections (movement from i to j facilitated by one or several other intermediate PAs). The maximum product probability (p*_ij_) is calculated through network analysis using the values of the direct dispersal probabilities between nodes (p_ij_). It holds that p*_ij_ ≥ p_ij_. Both probabilities will be equal when the direct movement is the most favorable (probable) pathway between i and j. p*_ij_ will be larger than p_ij_ when intermediate stepping stones increase the connectivity between i and j beyond what possible by using only the direct connection between them (*42,43,44*). In this analysis, p_ij_=1 when protected area i and j are connected (edge to edge) by a continuous pathway of intact land and p_ij_=0 if not. Two protected areas may not be directly connected by intact lands (hence having p_ij_=0), but may be connected through an intermediate stepping-stone protected areas, which would give p*_ij_=1.

### Connectivity between protected areas

The Probability of Connectivity (PC) metric accounts for both intra-PA (i=j) and inter-PA area (i≠j) connectivity, which is respectively given by the intra-PA (PCintra) and inter-PA (PCinter) components of PC as follows:

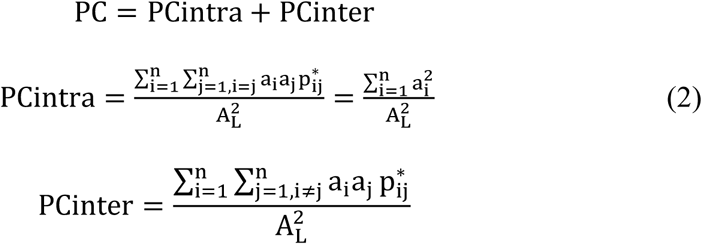

We focused on the inter-PA connectivity, given by PCinter. PCinter is defined as the probability that two points randomly located in two different PAs within the study area (therefore considering only the cases where i≠j) are connected to each other via intact habitat. We calculated PCinter using two scenarios: PCinter_ intact and PCinter_all. PCinter_intact is the value when considering that only the intact lands provide connectivity between PAs. PCinter_all is the value when any land (all land, intact, or not) provides connectivity between PAs, i.e. considering that two protected areas are connected when they are located in the same land mass or island. This analysis (PCinter_all) provided us with the maximum terrestrial protected area connectivity that could be theoretically achieved in a country or continent if all of its land was intact. In both scenarios, an 8-neighbouhood rule between land cells was used when defining the continuity of land (using the 1 km^2^ resolution of the human footprint layer).

### Connectivity provided by intact lands: ConnIntact

We combined PCinter_intact and PCinter_all, as defined above, to obtain ConnIntact, which quantifies the percentage of the protected are system that is connected through intact pathways. It is calculated using the following ratio:

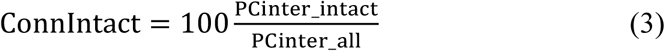

Which, given the equation for PCinter above, can be expressed as:

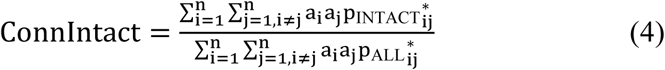

where p_INTACT_ refers to the maximum product probabilities when only the intact lands provide connectivity between protected areas, and p_ALL_ refers to the maximum product probabilities when all land would be intact and hence would provide the highest possible connectivity between protected areas. As noted above, n is the total number of protected areas in the study area (e.g. a country or continent), and a_i_ is the attribute of protected area i. ConnIntact provides the percentage of the protected area network that is connected by intact lands. This metric is expressed as percentage of the total area under protection.

## Acknowledgements

This research was supported by an Australian Government Research Training Program Scholarship and by the institutional activities of the Joint Research Centre of the European Commission;

## Author contributions

J.E.M.W conceived the study, J.E.M.W, O.V, S.S, M.W, B.W, J.A, and N.D developed the idea and methods, S.S and G.D developed and globally applied the protected area connectivity model, M.W. conducted the analysis of results and statistical evaluation, and M.W. led the writing of the manuscript with input from all authors.

## Data and materials availability

Data and materials availability: The human footprint data are available for download from https://datadryad.org/stash/dataset/doi:10.5061/dryad.052q5, protected-area data (2019 version) are available for download from www.protectedplanet.net by request. All other data needed to evaluate the conclusions in the paper are present in the paper or the supplementary materials.

## Supplementary Materials

Materials and Methods

Table S1 – S3

Fig S1 – S7

