## Supplementary data for "Only ten percent of the global terrestrial protected area network is connected via intact land"

**This PDF file includes:**

Materials and Methods

Supplementary Text

Figs S1 to S7

Tables S1 to S3

**Supplementary Text:**

Sensitivity analysis

We tested the sensitivity of our results to the data obtained from the human footprint score <4, using two additional human footprint thresholds. With a human footprint score <1, globally the proportion of connected protected areas results in 8.2% rather than 9.7% for a human footprint threshold of <4 (1.5% when considering the absolute difference in the proportion of connected protected areas globally). When a human footprint threshold of <10 is considered instead of <4, the proportion of the area under protection that is connected increases from 9.7% to 43.5% globally. This result does not alter our conclusions appreciably because the percentage of total land considered intact also varies in a similar fashion: 25% of all terrestrial land under <1 HFP threshold, 41.6% of all terrestrial land under <4 HFP threshold, and 74% of all terrestrial land under <10 HFP threshold.

Certain areas change considerably under different human footprint thresholds. For example, the increase in the protected area connectivity for human footprint threshold <10 is particularly noticeable in Oceania (from 15.6% to 94.9%) and Russia (from 1.9% to 79%), and raises to more than 30% of protected areas connected in Africa, Americas and Asia. This occurs because for this human footprint threshold, most of the roads are no longer considered as a barrier, which can influence the movement of some particular species or group of species *(11)*. This suggest that the connectivity depends also on how species respond to the permeability of the landscape. Therefore, the human footprint threshold of <1 might be better to assess the connectivity for sensitive species to human activities such as the boreal woodland caribou, while for species that can move through more human-modified landscapes, such as the American black bear, the human footprint threshold of <10 would make more sense.

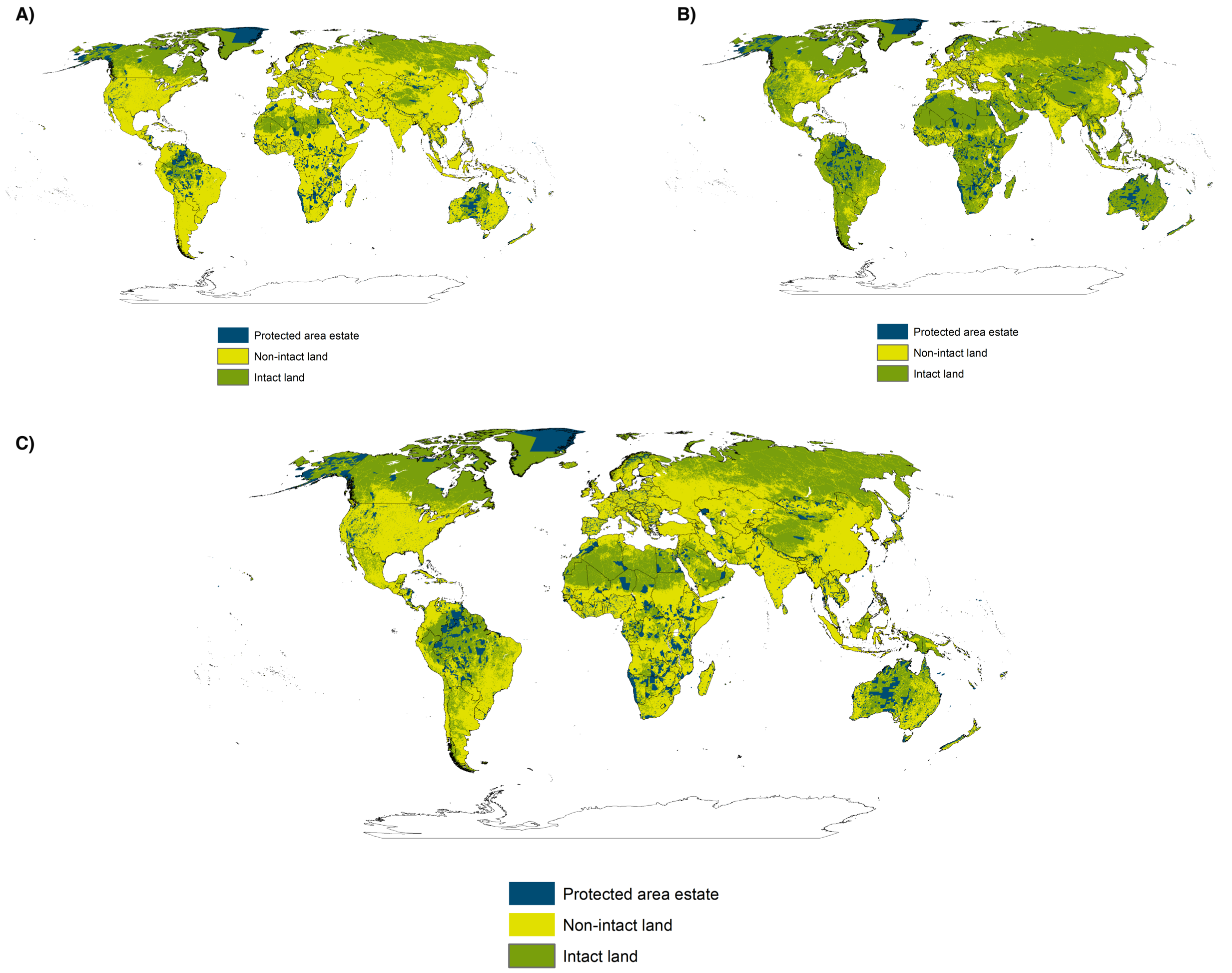


Fig. S1. Protected area estate. Global non-intact land (yellow), intact land (green), and the spatial distribution of protected area estate (blue) for A) human footprint value <1, B) human footprint value <10, and C) human footprint value <4.

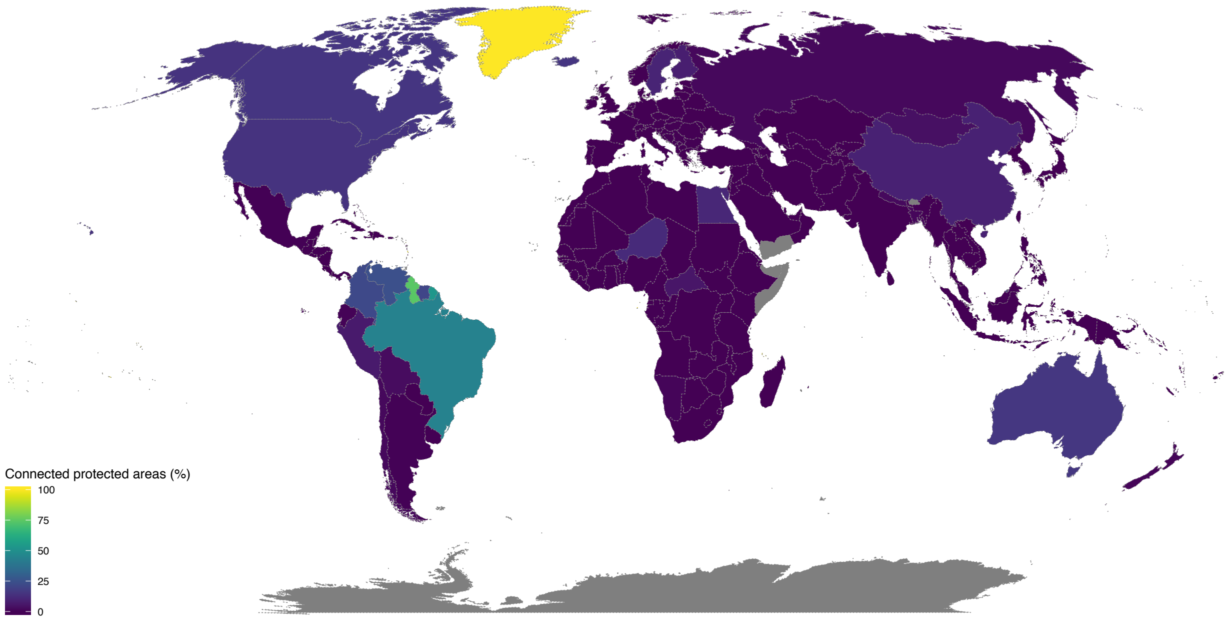


Fig. S2. Global map showing the proportion of protected areas connected through intact landscapes. Protected areas are regarded as connected if joined by areas with a human footprint value <1.

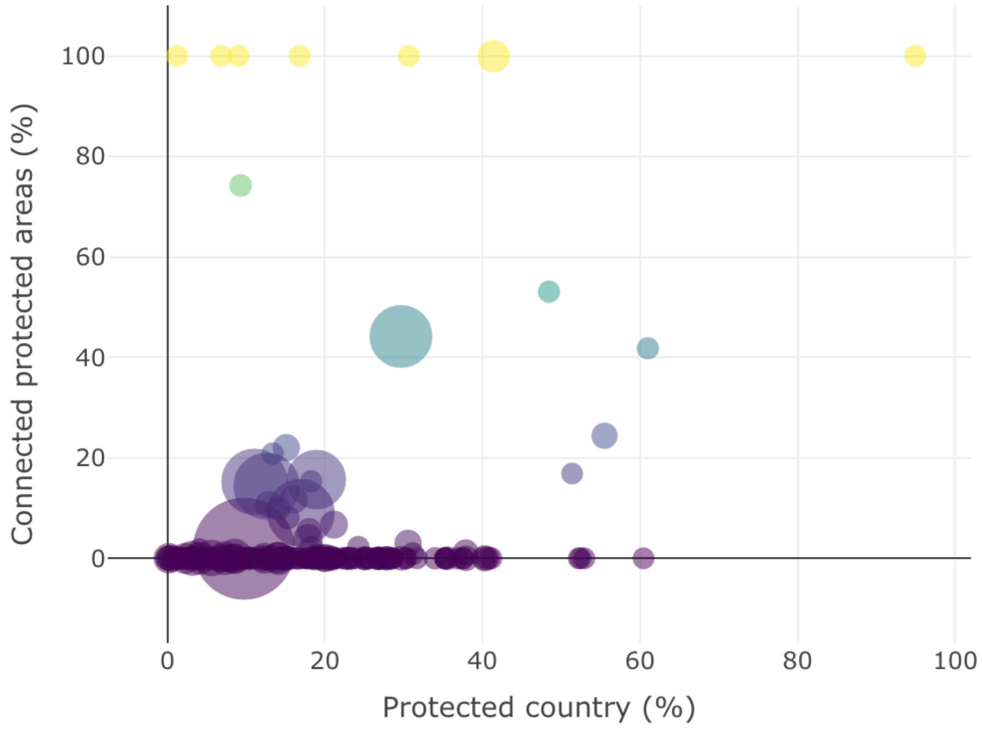


Fig. S3. Scatterplot showing the relationship between the proportion of land under protection and the proportion of connected protected areas per country. The size of the bubble highlighting the size of the country. Protected areas are regarded as connected if joined by areas with a human footprint value <1.

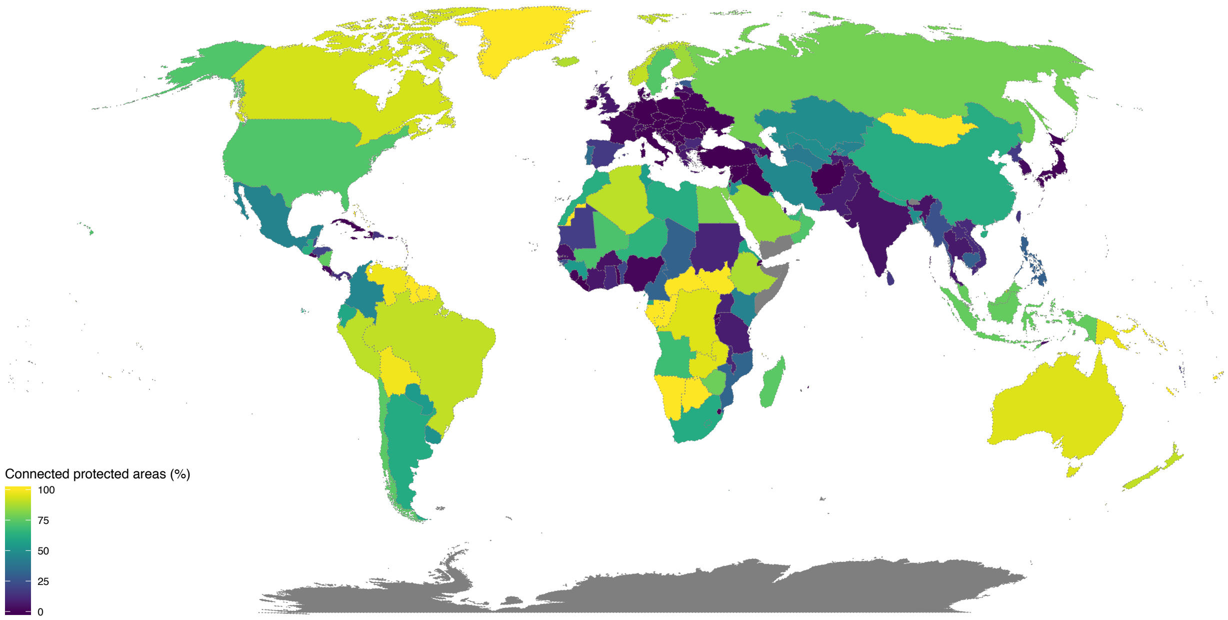


Fig. S4. Global map showing the proportion of protected areas connected through intact landscapes. Protected areas are regarded as connected if joined by areas with a human footprint value <10.

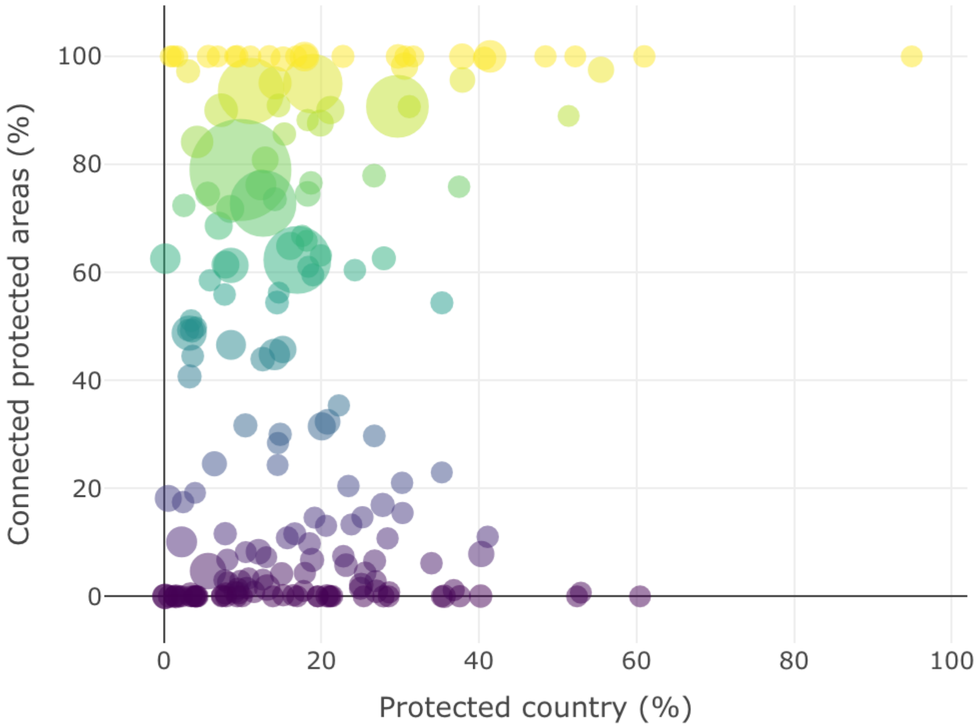


Fig. S5. Scatterplot showing the relationship between the proportion of land under protection and the proportion of connected protected areas per country. The size of the bubble highlighting the size of the country. Protected areas are regarded as connected if joined by areas with a human footprint value <10.

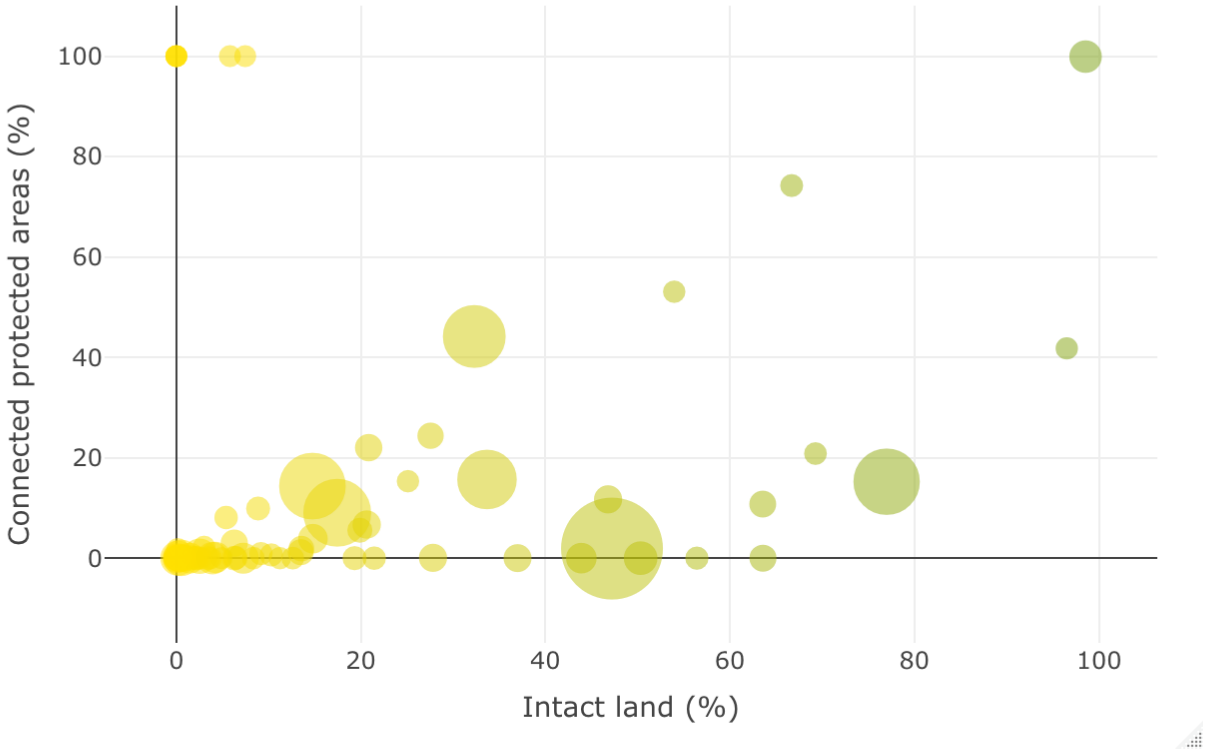


Fig. S6. Scatterplot showing the relationship between proportion of intact land and proportion of connected protected areas per country. The size of the bubble highlighting the size of the country. Protected areas are regarded as connected if joined by areas with a human footprint value <1.

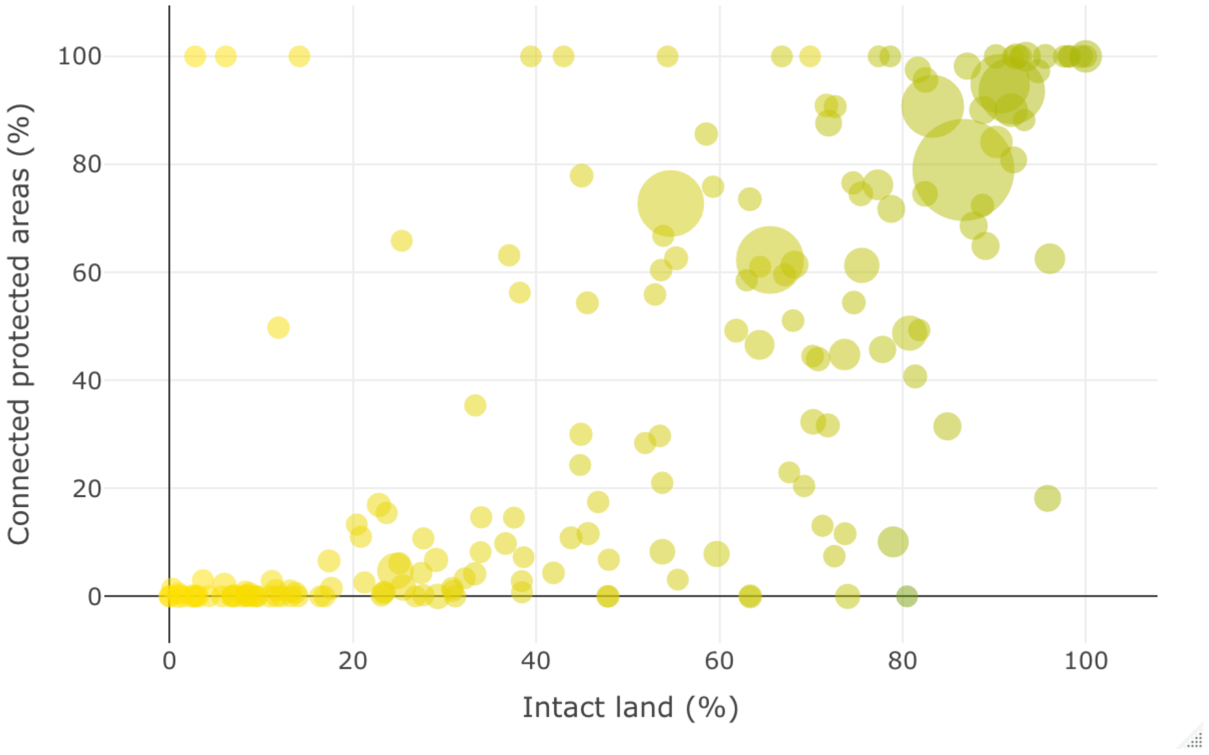


Fig. S7. Scatterplot showing the relationship between proportion of intact land and proportion of connected protected areas per country. The size of the bubble highlighting the size of the country. Protected areas are regarded as connected if joined by areas with a human footprint value <10.

Table S1. Sensitivity analysis of human footprint (HFP) thresholds showing the proportion of protected areas connected through intact pathways (ConnIntact %). All sensitivity analyses were performed considering those PAs with a size of at least 10 $km^2$.

| | Connected protected areas (%) | | |
|---|---|---|---|
| Areas of study | ConnIntact (%) HFP <1 | ConnIntact (%) HFP <4 | ConnIntact (%) HFP <10 |
| Africa | 0.1 | 0.5 | 34.8 |
| Americas | 12.7 | 14.8 | 44.4 |
| Asia | 2.2 | 3.2 | 42.2 |
| Europe | 0.1 | 0.3 | 2.3 |
| Oceania | 15.6 | 16.8 | 94.9 |
| Russia* | 1.9 | 4.5 | 79.0 |
| Global | 8.2 | 9.7 | 43.5 |

*Russia was treated as a separate entity because of its large size and large amount of connected land.

Table S2. Sensitivity analysis showing the proportion of all land that is intact under different human footprint (HFP) thresholds.

| | **Percentage of intact land** | | |
|---|---|---|---|
| | HFP <1 | HFP <4 | HFP <10 |
| **Global** | 25.0% | 41.6% | 74.0% |

Table S3. Country-level analysis of human footprint (HFP) key results showing the proportion of protected areas connected through intact lands (using HFP <1, <4, and <10).

| Country | Number PAs | Area PAs (km2) | ConnIntact % HFP <1 | ConnIntact % HFP <4 | ConnIntact_% HFP <10 |
|---|---|---|---|---|---|
| "AFG" | 2 | 669.8 | 0.0 | 0.0 | 0.0 |
| "AGO" | 13 | 86912.2 | 0.0 | 0.0 | 68.6 |
| "ALB" | 23 | 4859.6 | 0.0 | 0.0 | 1.0 |
| "AND" | 4 | 119.2 | 0.0 | 0.0 | 0.0 |
| "ARE" | 23 | 12475.6 | 0.0 | 0.0 | 66.8 |
| "ARG" | 220 | 233509.8 | 0.4 | 1.4 | 61.3 |
| "ARM" | 19 | 6778.5 | 0.0 | 0.0 | 13.3 |
| "ATG" | 2 | 30.0 | 100.0 | 100.0 | 100.0 |
| "AUS" | 2284 | 1454462.1 | 15.7 | 16.8 | 94.9 |
| "AUT" | 91 | 23007.6 | 0.0 | 0.0 | 0.0 |
| "AZE" | 27 | 8703.2 | 0.0 | 0.0 | 1.6 |
| "BDI" | 12 | 1974.7 | 0.0 | 3.0 | 3.0 |
| "BEL" | 52 | 5913.5 | 0.0 | 0.0 | 0.0 |
| "BEN" | 29 | 34099.5 | 0.0 | 0.0 | 21.0 |
| "BFA" | 49 | 40929.7 | 0.0 | 0.0 | 4.2 |
| "BGD" | 53 | 5290.1 | 1.8 | 1.8 | 49.8 |
| "BGR" | 77 | 44625.7 | 0.0 | 0.0 | 11.0 |
| "BHS" | 16 | 3170.8 | 0.0 | 68.9 | 100.0 |
| "BIH" | 4 | 692.7 | 0.0 | 0.0 | 0.0 |
| "BLR" | 101 | 19225.4 | 0.0 | 0.0 | 0.6 |
| "BLZ" | 19 | 8048.3 | 0.0 | 0.0 | 23.0 |

| | | | | | |
|---|---|---|---|---|---|
| "BOL" | 80 | 331003.7 | 3.0 | 16.3 | 98.2 |
| "BRA" | 774 | 2478601.9 | 44.2 | 51.5 | 90.7 |
| "BRN" | 8 | 2707.8 | 16.9 | 89.0 | 89.0 |
| "BWA" | 8 | 168479.1 | 0.0 | 0.0 | 100.0 |
| "CAF" | 16 | 112057.3 | 5.6 | 5.6 | 100.0 |
| "CAN" | 1576 | 1006607.5 | 15.2 | 23.3 | 93.6 |
| "CHE" | 44 | 3226.0 | 0.0 | 0.0 | 2.6 |
| "CHL" | 311 | 135812.5 | 1.9 | 30.3 | 74.5 |
| "CHN" | 1456 | 1591167.4 | 9.0 | 12.9 | 62.3 |
| "CIV" | 195 | 73541.9 | 0.0 | 0.0 | 5.8 |
| "CMR" | 34 | 48964.4 | 0.0 | 0.0 | 31.7 |
| "COD" | 30 | 320185.1 | 0.0 | 0.0 | 95.1 |
| "COG" | 13 | 139015.8 | 0.0 | 39.2 | 99.6 |
| "COL" | 192 | 167607.2 | 22.0 | 24.9 | 45.7 |
| "COM" | 3 | 168.5 | 100.0 | 100.0 | 100.0 |
| "CRI" | 30 | 13726.0 | 0.0 | 2.5 | 2.8 |
| "CUB" | 59 | 15762.1 | 0.0 | 0.0 | 0.2 |
| "CUW" | 2 | 44.1 | 0.0 | 0.0 | 0.0 |
| "CYP" | 9 | 3259.7 | 0.0 | 0.0 | 0.0 |
| "CZE" | 64 | 16506.1 | 0.0 | 0.0 | 0.0 |
| "DEU" | 715 | 124260.0 | 0.0 | 0.0 | 0.0 |
| "DJI" | 3 | 337.4 | 0.0 | 0.0 | 0.0 |
| "DMA" | 2 | 158.9 | 0.0 | 0.0 | 0.0 |
| "DNK" | 75 | 3292.6 | 0.0 | 0.0 | 0.5 |
| "DOM" | 57 | 12182.2 | 0.0 | 0.0 | 14.6 |
| "DZA" | 42 | 173376.5 | 0.0 | 0.0 | 90.0 |
| "ECU" | 50 | 46984.4 | 0.6 | 5.1 | 59.5 |

| | | | | | |
|---|---|---|---|---|---|
| "EGY" | 21 | 128104.9 | 10.8 | 10.8 | 80.8 |
| "ERI" | 3 | 5903.4 | 0.0 | 0.0 | 58.5 |
| "ESH" | 2 | 15047.6 | 0.0 | 0.0 | 100.0 |
| "ESP" | 458 | 138779.8 | 0.0 | 0.3 | 16.9 |
| "EST" | 102 | 6300.3 | 0.0 | 0.0 | 28.4 |
| "ETH" | 21 | 198744.5 | 0.0 | 0.4 | 87.6 |
| "FIN" | 287 | 46508.9 | 8.1 | 10.1 | 85.6 |
| "FJI" | 3 | 314.4 | 0.0 | 0.0 | 100.0 |
| "FRA" | 339 | 137229.9 | 0.0 | 0.0 | 2.1 |
| "GAB" | 23 | 58596.9 | 0.0 | 6.4 | 100.0 |
| "GBR" | 246 | 64738.6 | 0.0 | 0.7 | 6.6 |
| "GEO" | 37 | 5611.6 | 0.0 | 0.0 | 6.8 |
| "GHA" | 146 | 35668.5 | 0.0 | 0.0 | 10.8 |
| "GIN" | 63 | 86726.0 | 0.0 | 0.0 | 54.4 |
| "GLP" | 2 | 1075.9 | 0.0 | 0.0 | 0.0 |
| "GMB" | 4 | 399.5 | 0.0 | 0.0 | 0.0 |
| "GNB" | 23 | 4061.8 | 0.0 | 0.0 | 24.3 |
| "GNQ" | 10 | 5143.5 | 0.0 | 0.0 | 61.0 |
| "GRC" | 297 | 43791.6 | 0.0 | 0.0 | 6.1 |
| "GRL" | 85 | 897168.2 | 99.9 | 100.0 | 100.0 |
| "GTM" | 49 | 21364.7 | 0.0 | 0.0 | 63.1 |
| "GUF" | 7 | 44081.0 | 53.1 | 99.6 | 100.0 |
| "GUM" | 2 | 105.5 | 0.0 | 0.0 | 0.0 |
| "GUY" | 6 | 18323.7 | 74.2 | 100.0 | 100.0 |
| "HND" | 59 | 26220.8 | 0.0 | 0.0 | 20.4 |
| "HRV" | 63 | 20592.3 | 0.0 | 0.0 | 1.2 |
| "HTI" | 4 | 530.6 | 0.0 | 0.0 | 0.0 |

| | | | | | |
|---|---|---|---|---|---|
| "HUN" | 169 | 19006.1 | 0.0 | 0.0 | 0.0 |
| "IDN" | 367 | 223312.5 | 0.0 | 24.3 | 76.2 |
| "IND" | 417 | 166953.2 | 0.1 | 0.1 | 4.7 |
| "IRL" | 82 | 7892.3 | 0.0 | 0.1 | 0.8 |
| "IRN" | 107 | 138957.6 | 0.0 | 3.9 | 46.6 |
| "IRQ" | 4 | 6670.4 | 0.0 | 0.0 | 0.0 |
| "ISL" | 18 | 18301.2 | 15.3 | 15.8 | 88.2 |
| "ISR" | 25 | 3926.0 | 0.0 | 0.0 | 65.8 |
| "ITA" | 490 | 60933.2 | 0.0 | 0.0 | 0.1 |
| "JAM" | 8 | 1499.4 | 0.0 | 0.0 | 0.0 |
| "JOR" | 12 | 2742.1 | 0.0 | 0.8 | 49.3 |
| "JPN" | 531 | 98313.3 | 0.0 | 0.0 | 0.9 |
| "KAZ" | 75 | 86893.4 | 0.0 | 0.0 | 48.8 |
| "KEN" | 108 | 71457.3 | 0.0 | 0.0 | 43.9 |
| "KGZ" | 38 | 7037.1 | 0.0 | 0.0 | 44.5 |
| "KHM" | 25 | 47163.2 | 0.0 | 3.9 | 29.7 |
| "KIR" | 2 | 136.0 | 100.0 | 100.0 | 100.0 |
| "KOR" | 96 | 9474.7 | 0.0 | 0.0 | 2.8 |
| "KWT" | 11 | 2601.7 | 0.0 | 0.0 | 56.3 |
| "LAO" | 20 | 38363.5 | 0.0 | 0.0 | 11.6 |
| "LBN" | 4 | 232.8 | 0.0 | 0.0 | 0.0 |
| "LBR" | 6 | 3971.4 | 0.0 | 0.0 | 0.0 |
| "LBY" | 6 | 3399.4 | 0.0 | 0.0 | 62.5 |
| "LCA" | 2 | 103.3 | 0.0 | 0.0 | 0.0 |
| "LKA" | 80 | 19003.4 | 0.0 | 0.0 | 15.4 |
| "LTU" | 82 | 10300.0 | 0.0 | 0.0 | 0.2 |
| "LUX" | 11 | 1274.5 | 0.0 | 0.0 | 0.0 |

| | | | | | |
|---|---|---|---|---|---|
| "LVA" | 72 | 11130.2 | 0.0 | 0.0 | 4.3 |
| "MAR" | 137 | 124646.0 | 0.0 | 2.8 | 62.6 |
| "MDA" | 12 | 1275.3 | 0.0 | 0.0 | 0.0 |
| "MDG" | 69 | 32400.8 | 0.0 | 0.0 | 74.5 |
| "MEX" | 378 | 272884.5 | 0.2 | 1.5 | 44.8 |
| "MKD" | 11 | 2624.3 | 0.0 | 0.0 | 8.2 |
| "MLI" | 18 | 102817.9 | 0.0 | 3.8 | 71.8 |
| "MMR" | 36 | 41959.5 | 0.0 | 0.0 | 24.6 |
| "MNE" | 7 | 1447.0 | 0.0 | 0.0 | 3.3 |
| "MNG" | 95 | 277555.3 | 3.9 | 4.0 | 100.0 |
| "MOZ" | 24 | 163469.5 | 0.0 | 0.0 | 32.3 |
| "MRT" | 5 | 5833.9 | 0.0 | 0.0 | 18.2 |
| "MTQ" | 2 | 1044.2 | 100.0 | 100.0 | 100.0 |
| "MUS" | 2 | 71.5 | 0.0 | 0.0 | 0.0 |
| "MWI" | 44 | 26776.2 | 0.0 | 0.2 | 10.7 |
| "MYS" | 236 | 61370.2 | 0.2 | 11.4 | 76.6 |
| "NAM" | 37 | 311782.4 | 1.2 | 5.3 | 100.0 |
| "NCL" | 12 | 9546.2 | 0.0 | 21.2 | 100.0 |
| "NER" | 12 | 203268.7 | 11.7 | 11.7 | 64.9 |
| "NGA" | 517 | 119354.7 | 0.0 | 0.0 | 1.6 |
| "NIC" | 49 | 45102.3 | 0.0 | 0.0 | 75.9 |
| "NLD" | 72 | 3161.7 | 0.0 | 0.1 | 1.4 |
| "NOR" | 194 | 53197.0 | 0.0 | 0.3 | 90.9 |
| "NPL" | 17 | 34763.1 | 2.2 | 2.2 | 60.4 |
| "NZL" | 382 | 82026.2 | 0.9 | 6.3 | 90.7 |
| "OMN" | 6 | 7895.0 | 0.0 | 0.0 | 72.4 |
| "PAK" | 112 | 92705.4 | 0.0 | 0.1 | 8.3 |

| | | | | | |
|---|---|---|---|---|---|
| "PAN" | 24 | 15311.4 | 0.0 | 2.2 | 13.1 |
| "PER" | 99 | 270613.8 | 6.7 | 58.8 | 90.1 |
| "PHL" | 107 | 44042.7 | 0.0 | 14.3 | 30.0 |
| "PLW" | 2 | 50.6 | 0.0 | 0.0 | 100.0 |
| "PNG" | 32 | 13997.5 | 0.0 | 16.0 | 97.3 |
| "POL" | 176 | 123138.0 | 0.0 | 0.0 | 0.1 |
| "PRI" | 12 | 382.0 | 0.0 | 0.0 | 0.0 |
| "PRK" | 13 | 2945.9 | 0.0 | 0.0 | 17.5 |
| "PRT" | 57 | 20356.6 | 0.0 | 0.0 | 35.4 |
| "PRY" | 53 | 57091.7 | 0.0 | 0.1 | 54.4 |
| "PSE" | 17 | 442.7 | 0.0 | 0.0 | 0.0 |
| "PYF" | 2 | 44.3 | 100.0 | 100.0 | 100.0 |
| "QAT" | 3 | 1510.2 | 0.0 | 0.0 | 7.2 |
| "ROU" | 200 | 57430.0 | 0.0 | 0.0 | 1.5 |
| "RUS" | 2001 | 1593180.8 | 1.9 | 4.5 | 79.0 |
| "RWA" | 6 | 2297.2 | 0.0 | 0.0 | 0.0 |
| "SAU" | 26 | 90849.3 | 0.0 | 0.0 | 84.2 |
| "SDN" | 10 | 42083.2 | 0.0 | 0.0 | 10.1 |
| "SEN" | 90 | 49199.7 | 0.0 | 0.0 | 4.3 |
| "SJM" | 24 | 37843.8 | 41.8 | 100.0 | 100.0 |
| "SLB" | 5 | 247.7 | 0.0 | 100.0 | 100.0 |
| "SLE" | 30 | 6507.2 | 0.0 | 0.0 | 0.8 |
| "SLV" | 9 | 1638.9 | 0.0 | 0.0 | 0.0 |
| "SRB" | 44 | 6477.4 | 0.0 | 0.0 | 0.1 |
| "SSD" | 15 | 97578.0 | 0.0 | 0.0 | 99.5 |
| "STP" | 2 | 294.3 | 100.0 | 100.0 | 100.0 |
| "SUR" | 10 | 20863.0 | 20.8 | 52.8 | 100.0 |

| | | | | | |
|---|---|---|---|---|---|
| "SVK" | 41 | 18062.1 | 0.0 | 0.0 | 0.0 |
| "SVN" | 27 | 10661.6 | 0.0 | 0.0 | 0.7 |
| "SWE" | 293 | 57416.8 | 9.9 | 24.4 | 73.5 |
| "SWZ" | 5 | 711.5 | 0.0 | 0.0 | 0.0 |
| "SYR" | 5 | 1262.8 | 0.0 | 0.0 | 0.0 |
| "TCD" | 13 | 252495.6 | 0.1 | 10.2 | 31.5 |
| "TGO" | 28 | 15560.2 | 0.0 | 0.0 | 0.7 |
| "THA" | 131 | 96160.1 | 0.0 | 0.1 | 6.7 |
| "TJK" | 20 | 31632.4 | 0.0 | 0.6 | 7.4 |
| "TKM" | 19 | 15365.9 | 0.0 | 0.0 | 40.7 |
| "TLS" | 20 | 1871.0 | 0.0 | 0.1 | 3.1 |
| "TTO" | 14 | 1463.9 | 0.0 | 0.0 | 0.0 |
| "TUN" | 44 | 11986.9 | 0.0 | 9.2 | 55.9 |
| "TUR" | 14 | 1543.8 | 0.0 | 0.0 | 0.0 |
| "TWN" | 9 | 6925.1 | 0.0 | 0.0 | 14.6 |
| "TZA" | 234 | 356938.3 | 0.0 | 2.9 | 7.8 |
| "UGA" | 142 | 37079.0 | 0.0 | 0.0 | 9.8 |
| "UKR" | 242 | 19615.4 | 0.0 | 0.0 | 0.3 |
| "URY" | 17 | 6078.8 | 0.0 | 0.0 | 51.1 |
| "USA" | 3673 | 1153291.0 | 14.4 | 14.7 | 72.8 |
| "UZB" | 15 | 15169.1 | 0.0 | 0.0 | 49.2 |
| "VEN" | 78 | 489479.2 | 24.4 | 25.4 | 97.5 |
| "VNM" | 109 | 24175.0 | 0.0 | 0.0 | 11.6 |
| "VUT" | 8 | 483.3 | 0.0 | 0.0 | 19.2 |
| "ZAF" | 627 | 94936.3 | 0.0 | 0.1 | 61.4 |
| "ZMB" | 270 | 281763.5 | 0.0 | 0.3 | 95.6 |
| "ZWE" | 44 | 103261.5 | 0.0 | 1.2 | 77.9 |

11. M. A. Tucker, K. Böhning-Gaese, W. F. Fagan, J. M. Fryxell, B. Van Moorter, S. C. Alberts, A. H. Ali, A. M. Allen, N. Attias, T. Avgar, H. Bartlam-Brooks, B. Bayarbaatar, J. L. Belant, A. Bertassoni, D. Beyer, L. Bidner, F. M. van Beest, S. Blake, N. Blaum, C. Bracis, D. Brown, P. J. N. de Bruyn, F. Cagnacci, J. M. Calabrese, C. Camilo-Alves, S. Chamaillé-Jammes, A. Chiaradia, S. C. Davidson, T. Dennis, S. DeStefano, D. Diefenbach, I. Douglas-Hamilton, J. Fennessy, C. Fichtel, W. Fiedler, C. Fischer, I. Fischhoff, C. H. Fleming, A. T. Ford, S. A. Fritz, B. Gehr, J. R. Goheen, E. Gurarie, M. Hebblewhite, M. Heurich, A. J. M. Hewison, C. Hof, E. Hurme, L. A. Isbell, R. Janssen, F. Jeltsch, P. Kaczensky, A. Kane, P. M. Kappeler, M. Kauffman, R. Kays, D. Kimuyu, F. Koch, B. Kranstauber, S. LaPoint, P. Leimgruber, J. D. C. Linnell, P. López-López, A. C. Markham, J. Mattisson, E. P. Medici, U. Mellone, E. Merrill, G. de Miranda Mourão, R. G. Morato, N. Morellet, T. A. Morrison, S. L. D’iaz-Muñoz, A. Mysterud, D. Nandintsetseg, R. Nathan, A. Niamir, J. Odden, R. B. O'Hara, L. G. R.

Oliveira-Santos, K. A. Olson, B. D. Patterson, R. de Paula, L. Pedrotti, B. Reineking, M. Rimmler, T. L. Rogers, C. M. Rolandsen, C. S. Rosenberry, D. I. Rubenstein, K. Safi, S. Sa'id, N. Sapir, H. Sawyer, N. M. Schmidt, N. Selva, A. Sergiel, E. Shiilegdamba, J. P. Silva, N. Singh, E. J. Solberg, O. Spiegel, O. Strand, S. Sundaresan, W. Ullmann, U. Voigt, J. Wall, D. Wattles, M. Wikelski, C. C. Wilmers, J. W. Wilson, G. Wittemyer, F. Zik eba, T. Zwijacz-Kozica, T. Mueller, Moving in the Anthropocene: Global reductions in terrestrial mammalian movements. *Science.* **359**, 466–469 (2018).

24. IUCN, Increase connectivity between protected areas: A key solution to conserve biodiversity in Vietnam (2016), (International Union for Conservation of Nature, Switzerland, 2008).

25. A. J. Lynam, Securing a future for wild Indochinese tigers: Transforming tiger vacuums into tiger source sites. *Integr. Zool.* **5**, 324–334 (2010).

26. R. Stone, The Saola's Last Stand. *Nature*. **314**, 1380–1383 (2006).

27. C. Schwitzer, R. A. Mittermeier, S. E. Johnson, G. Donati, M. Irwin, H. Peacock, J. Ratsimbazafy, J. Razafindramanana, E. E. Louis, L. Chikhi, I. C. Colquhoun, J. Tinsman, R. Dolch, M. Lafleur, S. Nash, E. Patel, B. Randrianambinina, T. Rasolofoharivelo, P. C. Wright, Conservation. Averting lemur extinctions amid Madagascar's political crisis. *Science*. **343**, 842–843 (2014).

28. S. H. M. Butchart, M. Clarke, R. J. Smith, R. E. Sykes, J. P. W. Scharlemann, M. Harfoot, G. M. Buchanan, A. Angulo, A. Balmford, B. Bertzky, T. M. Brooks, K. E. Carpenter, M. T. Comeros-Raynal, J. Cornell, G. F. Ficetola, L. D. C. Fishpool, R. A. Fuller, J. Geldmann, H. Harwell, C. Hilton-Taylor, M. Hoffmann, A. Joolia, L. Joppa, N. Kingston, I. May, A. Milam, B. Polidoro, G. Ralph, N. Richman, C. Rondinini, D. B. Segan, B. Skolnik, M. D. Spalding, S. N. Stuart, A. Symes, J. Taylor, P. Visconti, J. E. M. Watson, L. Wood, N. D. Burgess, Shortfalls and Solutions for Meeting National and Global Conservation Area Targets. *Conserv. Lett.* **8**, 329–337 (2015).

29. A. Alfiky, G. Kaule, M. Salheen, Agricultural Fragmentation of the Nile Delta; A Modeling Approach to Measuring Agricultural Land Deterioration in Egyptian Nile Delta. *Procedia*

*Environ. Sci.* **14**, 79–97 (2012).

30. J. E. M. Watson, T. Evans, O. Venter, B. Williams, A. Tulloch, C. Stewart, I. Thompson, J. C. Ray, K. Murray, A. Salazar, C. McAlpine, P. Potapov, J. Walston, J. G. Robinson, M. Painter, D. Wilkie, C. Filardi, W. F. Laurance, R. A. Houghton, S. Maxwell, H. Grantham, C. Samper, S. Wang, L. Laestadius, R. K. Runting, G. A. Silva-Chávez, J. Ervin, D. Lindenmayer, The exceptional value of intact forest ecosystems. *Nat. Ecol. Evol.* **2**, 599–610 (2018).

31. J. M. Rey Benayas, A. C. Newton, A. Diaz, J. M. Bullock, Enhancement of biodiversity and ecosystem services by ecological restoration: a meta-analysis. *Science*. **325**, 1121–1124 (2009).

32. IUCN-WCPA, “Guidelines for Recognizing and ReporƟng Other Effective Area-based Conservation Measures: First Version,” (International Union for Conservation of Nature , Switzerland, 2017).

33. N. Dudley, H. Jonas, F. Nelson, J. Parrish, A. Pyhälä, S. Stolton, J. E. M. Watson, The essential role of other effective area-based conservation measures in achieving big bold conservation targets. *Glob. Ecol. Conserv.* **15**, e00424 (2018).

34. United Nations Environment Program, "UN Decade on Ecosystem Restoration (2018)," (United Nations, Switzerland, 2018).

35. R. Maggini, H. Kujala, M. Taylor, J. Lee, H. Possingham, B. Wintle, R. Fuller, “Protecting and restoring habitat to help Australia’s threatened species adapt to climate change,” (National Climate Change Adaptation Research Facility, 2013).

36. L. Cantú-Salazar, K. J. Gaston, Very Large Protected Areas and Their Contribution to

Terrestrial Biological Conservation. *Bioscience*. **60**, 808–818 (2010).

37. M. Schröter, R. Kraemer, S. Ceauşu, G. M. Rusch, Incorporating threat in hotspots and coldspots of biodiversity and ecosystem services. *Ambio*. **46**, 756–768 (2017).

38. J. E. M. Watson, M. C. Evans, J. Carwardine, R. A. Fuller, L. N. Joseph, D. B. Segan, M. F. J. Taylor, R. J. Fensham, H. P. Possingham, The Capacity of Australia's Protected-Area System to Represent Threatened Species. *Conserv. Biol.* **25**, 324–332 (2011).

39. C. N. Cook, R. S. Valkan, M. B. Mascia, M. A. McGeoch, Quantifying the extent of protected-area downgrading, downsizing, and degazettement in Australia. *Conserv. Biol.* **31**, 1039–1052 (2017).

40. M. S. Ward, J. S. Simmonds, A. E. Reside, J. E. M. Watson, J. R. Rhodes, H. P. Possingham, J. Trezise, R. Fletcher, L. File, M. Taylor, Lots of loss with little scrutiny: The attrition of habitat critical for threatened species in Australia. *Conserv. Sci. Pract.*, **1**, e117 (2019).

41. UN Environment World Conservation Monitoring Centre, International Union for Conservation of Nature, World Database on Protected Areas (2017); www.protectedplanet.net.

42. S. Saura, L. Pascual-Hortal, A new habitat availability index to integrate connectivity in landscape conservation planning: comparison with existing indices and application to a case study. *Landsc. Urban Plan*. **83**, 91–103 (2007).

43. S. Saura, L. A. Rubio, common currency for the different ways in which patches and links can contribute to habitat availability and connectivity in the landscape.

*Ecography*. **33**, 523–537 (2010).

44. S. Saura, Ö. Bodin, and M. J. Fortin, Stepping stones are crucial for species' long-distance dispersal and range expansion through habitat networks. *J Appl Ecol*, **51**, 171-182 (2014).
